# The holdase function of *Escherichia coli* Hsp70 (DnaK) chaperone

**DOI:** 10.1101/305854

**Authors:** Ricksen S. Winardhi, Qingnan Tang, Huijuan You, Michael Sheetz, Jie Yan

## Abstract

In *Escherichia coli*, the DnaK/DnaJ/GrpE system plays a critical role in mediating protein refolding and buffering against protein aggregation due to environmental stress. The underlying mechanism remains unclear. In this work, we probe the activity of DnaK/DnaJ/GrpE system with single-molecule protein refolding assay using tandem repeats of titin immunoglobulin 27 (I27)_8_. We provide direct evidence that DnaK in apo– and ADP-bound state is predominantly a holdase, which kinetically stabilizes the polyprotein in its unfolded form. Binding of ATP relieves DnaK’s holding, allowing protein refolding. The presence of co-chaperone DnaJ and GrpE modulates this holding-release switching, possibly by altering DnaK’s nucleotide state. Our findings thus provide important insights to the molecular mechanism of DnaK/DnaJ/GrpE system.

## INTRODUCTION

In virtually all living organisms, proteins are synthesized by a molecular machine called ribosome by sequential addition of amino acids that forms a polypeptide chain. The nascent polypeptide chain released from the ribosome then needs to fold into its native protein structure. This is a complex and vital process that often requires the assistance of other proteins, broadly termed as chaperones. There are many different families of chaperones, which are categorized according to their molecular weight, generally prefixed by the term ‘Hsp’ as historically these proteins are named ‘heat shock protein’ due to their high expression under stress condition such as heat shock [1].

Each of the chaperone family serves a different function in the general housekeeping of proteins in cells, and they often works in synergy with each other. For example, Hsp70 family of proteins assists client folding in its early stage by binding to the nascent polypeptide chain on the ribosomes, while Hsp90 assists during late stage folding. In addition, proteins that are damaged due to environmental stress need to be either repaired or degraded. Hsp60 can help to refold denatured proteins, as well as promote *de novo* folding of polypeptides. Another class of chaperones, Hsp100, utilizes energy from ATP to actively unfold and disaggregate misfolded protein structures and aggregates. In short, the complex network of chaperones serves many functions: assisting folding of newly synthesized or denatured proteins, preventing aggregation, unfolding of protein aggregates, all of which are crucial in order to maintain proteome integrity and protein homeostasis [2].

Among the various family of molecular chaperones Hsp70 is of special interest due to its implication in various neurodegenerative diseases, as many of these are caused by accumulation of aggregates/misfolded proteins [3]. Hsp70 also plays another critical role in regulating cell signaling, including type II diabetes and cancer-related signaling pathways [4]. Studies on Hsp70 system mainly focus on DnaK, which is the major bacterial Hsp70 in *E. coli*, as a model. DnaK comprises of two domains: N-terminal nucleotide-binding domain (NBD) and C-terminal substrate-binding domain (SBD), connected via a flexible and highly conserved linker [5]. The SBD contains a *β*-sheet peptide binding groove/cleft (SBD-*β*) for substrate binding, and an *α*-helical lid (SBD-*α*) for substrate locking.

The NBD is responsible for ATPase activity, which is important in switching the SBD conformation between open and closed state, i.e. whether the SBD-*α* is detached or attached to the SBD-*β*. In the apo-state and ADP-bound state, the NBD is undocked from the SBD, and the SBD-*α* is pressed against the SBD-*β* [5], while in the ATP-bound state the NBD is docked to the SBD with SBD-*α* separated away from SBD-*β* [6, 7]. Thus, ATP hydrolysis cycle dictates the conformational state of the SBD in DnaK to control binding activity to peptide substrate. Recently, the importance of SBD-*β* allostery in actively controlling substrate binding and release was elucidated [8]. As such, substrate affinity depends on whether the conformation of SBD is in the open or closed state. Substrate binding can also alter DnaK conformational state [9, 10], which has been suggested to stimulate ATPase activity observed upon substrate binding up to 25-fold [11–13]. Taken together, the substrate binding activity depends on the complex interplay of multiple factors: SBD-*β* dynamics, SBD-*α* lid opening/closure, peptide binding, and the nucleotide-state that tightly control allostericity.

In addition to its conformational dynamics, the activity of DnaK also depends on other factors, such as its oligomerization activity [14] and its interaction with other chaperones and co-chaperones [15]. In the ATP-bound state of DnaK (without peptide), the intrinsic ATPase activity of DnaK is very slow [0.02 min-1 at 25 ° C] [11, 16, 17]. Assistance of DnaJ, which is bacterial Hsp40 protein, can accelerate the rate ATP hydrolysis by a maximum of 15,000-fold at 5 °C [11], and induces the *α*-helical lid to close and lock the substrate in the SBD. Following ATP hydrolysis, a nucleotide exchange factor (NEF), GrpE, helps to facilitate nucleotide exchange by stimulating ADP release by 5,000-fold [18]. Together with its co-chaperones, DnaK forms the DnaK/DnaJ/GrpE system for efficient chaperoning activity in *E. coli*. DnaK also works in tandem with various other chaperones. For example, the DnaK is known to collaborate with HtpG (*E. coli* Hsp90) for client protein remodeling [19] and ClpB during protein disaggregation [20–22]. Importantly, DnaK serves as the central organizer in *E. coli* chaperone network to control proteostasis [23].

While many aspects of DnaK have been elucidated, questions remain on how DnaK binding to protein substrate may facilitate protein folding. Previously, DnaK was assumed to act as a holdase or an unfoldase [24, 25]. A holdase refers a chaperone binding to an unfolded substrate that decreases the rate of transition to the folded state. By binding to exposed hydrophobic region of unfolded or disaggregated protein segments, a holdase is expected to slow down the protein folding to prevent premature folding that may result in misfolding and/or aggregation. In contrast, an unfoldase refers to a chaperone that increases the unfolding rate of a folded/misfolded/aggregated substrate to facilitate the proteins to refold to their native conformation. The extent to which DnaK employ these mechanisms warrants further investigation.

In this article, we aim to address the above questions by using single-molecule protein refolding assay, which can provide a direct insight into how chaperone activity interferes directly with protein refolding. This method is based on magnetic tweezers that is capable of applying force to a protein domain and modulating the balance between the folded and the unfolded states [26]. This makes it possible to maintain the unfolded conformation at sufficiently high forces allowing chaperone binding, and switch to lower forces to investigate the effects of the chaperone on the protein refolding. We have recently demonstrated the use of this single-molecule assay in studying unfolding and refolding of several typical protein domains under force, such as Ig domains [27, 28], *α*-helix bundles [29, 30] and spectrin repeats [31] under force. We have also applied this approach to probe small molecule binding to unfolded polyprotein [32].

Compared to bulk biochemical assays, this label-free single-molecule manipulation assay has several advantages: 1) chaperone-substrate interaction and the effect on substrate refolding can be studied in the absence of inteference from the commonly used denaturants, 2) protein substrate can be reliably unfolded and refolded over a long period of time, and 3) single-protein sensitivity can be achieved. We find that DnaK in the apo– and ADP-bound state can serve as a very effective holdase. Upon binding of ATP, DnaK holdase activity is relieved to allow for protein refolding, while addition of DnaJ in the presence of ATP converts DnaK back to a holdase. We also find that further addition of GrpE can relieve this DnaJ-dependent holdase activity of DnaK in the presence of ATP.

## MATERIALS AND METHODS

### Expression and Purification of DnaK, DnaJ, and GrpE

DnaK, DnaJ, and GrpE proteins were obtained as follows. The *dnak, dnaj*, and *grpe* genes were each cloned into the pNIC28-Bsa4 vector [33]. BL21(DE3) Rosetta competent cells were transformed with each of the recombinant plasmids. The cells were cultivated in TB media supplemented with 100 *μ*g/mL kanamycin and 34 *μ*g/mL chloramphenicol in a LEX system (Harbinger Biotech). Overexpression of proteins was induced with 0.5 mM IPTG overnight at 18°C. The cells were then harvested by centrifugation and resuspended in lysis buffer (100 mM HEPES, 500 mM NaCl, 10 mM Imidazole, 10 % glycerol, 0.5 mM TCEP, pH 8.0) containing 1000 U benzonase (Merck) and 1000x dilution protease inhibitor cocktail (Calbiochem). Following this, the cells were sonicated and the lysates were centrifuged at 47,000 g, 4 °C. for 25 min. The supernatants were filtered through 1.2 *μ*m syringe filters and loaded onto IMAC columns in an AKTA Xpress system (GE Healthcare). Bound proteins were washed with buffers containing up to 25 mM imidazole and finally eluted with buffer containing 500 mM imidazole. Eluted proteins were loaded onto HiLoad 16/60 Superdex 200 columns (GE Healthcare) equilibrated with gel filtration buffer (20 mM HEPES pH 7.5, 300 mM NaCl, 10 % (v/v) glycerol, 2 mM TCEP) and eluted at 1.2 mL/min. Protein fractions were analyzed on SDS-PAGE gels; fractions containing target proteins were then pooled and concentrated with Vivaspin 20 centrifugal concentrators (VivaScience). The final protein concentration was assessed by measuring absorbance at 280 nm on Nanodrop ND-1000 (Nano-Drop Technologies). Purified proteins were aliquoted into smaller fractions, frozen in liquid nitrogen and stored at −80 °C‥

### Protein construct for single protein refolding assay

Protein construct of HaloTag-(I27)_8_-Avitag and SpyTag-(I27)_8_-Avitag was obtained by protein overex pression in *Escherichia coli* bacteria. The Avitag was biotinylated by co-expression *in vivo* with BirA enzyme. In wild type I27 domain, there are two cysteines at residues 47 and 63 [34]. To avoid potential formation of disulfide bond between the two cysteines, the two cysteines were substituted with alanines. These purified polyprotein construct was used as the substrate for binding in the single-protein refolding assay described below.

### Nucleotide

ATP, ATP*γ*s, and ADP were purchased from Sigma-Aldrich. For all experiments conducted in the presence of ATP, an additional regeneration system comprising 0.1 mg/mL pyruvate kinase (PK) from rabbit muscle and 2 mM phosphoenolpyruvate (PEP) were added. For all experiments conducted in the presence of ADP, the ADP solution used was treated with hexokinase and D-glucose to remove the contaminating ATP as described previously [35–37].

### Quantification of ATP contamination

ATP contamination in protein samples and nucleotide solution were quantified using ENLITEN ATP assay system (Promega). The protein samples were first heated at 95 ^¤^C for denaturation and nucleotide release. After the sample was brought to room temperature, regeneration system (see Nucleotide section above) was added to convert the ADP into ATP, such that the total amount of ATP and ADP contamination in the protein samples can be quantified.

### Single-protein refolding assay

Single-protein refolding experiments were carried out using in-house developed back-scattered vertical magnetic tweezers [26]. The extension change of the molecules was measured based on the change of the height of the end attached super-paramagnetic bead from the coverslip surface at a resolution of ∼ 1 nm [26, 38]. Force was calibrated based on the method described in [26, 38], which has a 10% relative error due to the uncertainty in the size of the super paramagnetic beads. In our experiments, tandem repeat of I27 immunoglobulin domain, (I27)_8_, was used as a model binding substrate for studying chaperone binding and its effect on protein refolding, because its mechanical stability has been well characterized [28]. To form the construct for protein stretching, (I27)_8_ polyprotein is tethered on one end to a glass coverslip surface using either HaloTag or SpyTag system [39, 40], while another end is tethered to a magnetic bead via streptavidin-biotin complex. This setup is contained within a flow chamber that allows convenient solution exchange.

Protein refolding was quantified by performing multiple cycles of protein unfolding-refolding experiments (Fig. 1). Each cycle consists of force-increase (to unfold proteins) and force-decrease scan (to allow for protein refolding). In the force-increase scan, the force is increased from 1 ± 0.1 pN to 80 ± 8 pN with a constant loading rate of 2 pN/s. The tether was held at 80 ± 8 or higher until all of the protein domains in the tether were unfolded. Next, the force was exponentially reduced to 1 ± 0.1 pN by increasing bead-magnet separation with a constant speed of 150 *μ*m/s. As I27 domain refolding largely occurs at forces < 5 pN, this force-decrease procedure allows for chaperone binding during the 12 s time window when force is reduced from 80 ± 8 pN to 5 ± 0.5 pN, while allowing protein refolding to take place during the 13 s time window when the force is reduced from 5 ± 0.5pN to 1 ± 0.1 pN. The tether was then held at 1 ± 0.1 pN for an additional 10 s before continuing to the next cycle. All the experiments were performed using this protocol, unless otherwise stated.

**FIG. 1.**
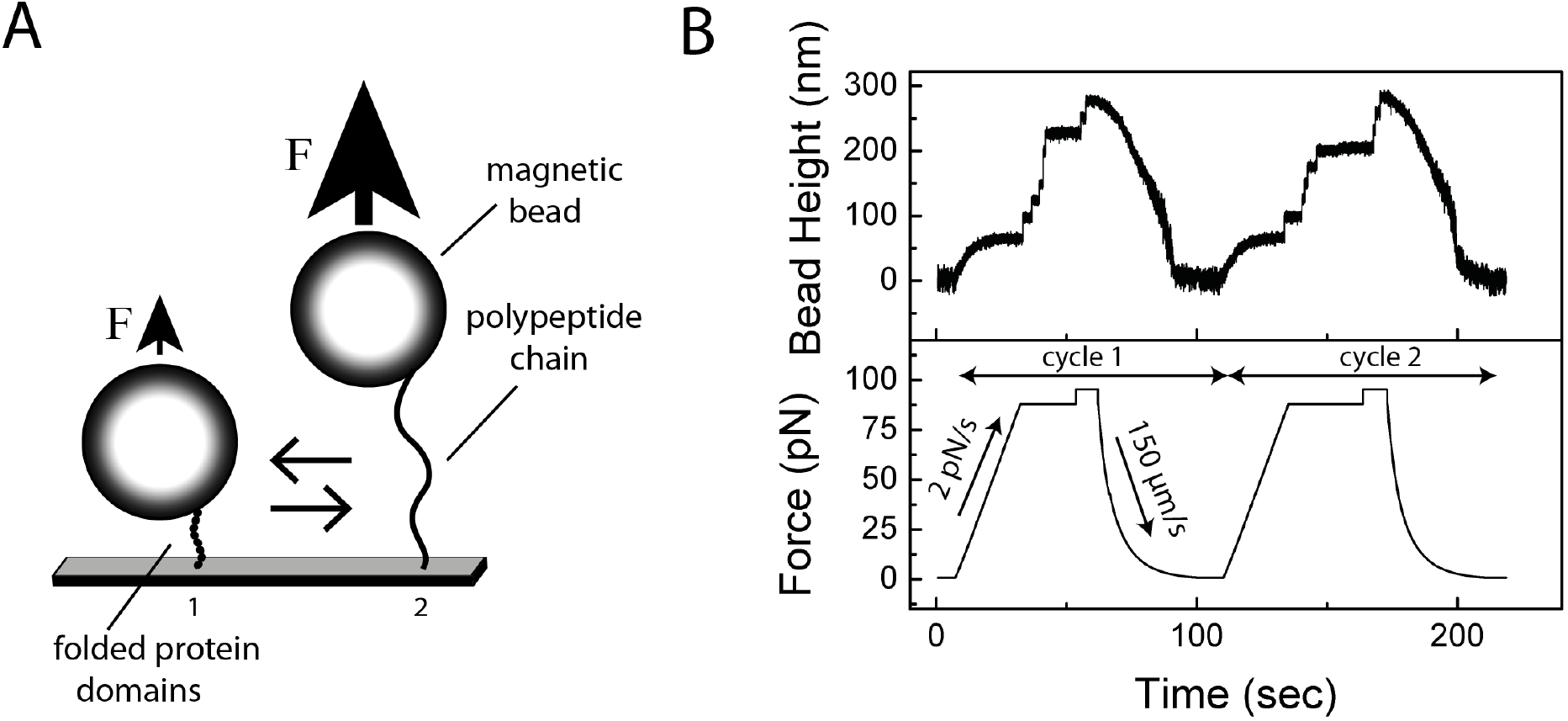
Mechanical unfolding and refolding of (I27)_8_ using magnetic tweezers. (A) Schematic of our experimental setup with protein substrate tethered in between glass coverslip and magnetic bead. A pair of permanent magnet is placed above the sample. Decreasing the distance between the magnet and the magnetic bead results in a higher force and hence unfolding of protein domains. Conversely, increasing the distance results in a lower force that allows protein to refold. This cycle of unfolding and refolding is repeated many times and the fraction of protein refolding is obtained. (B) A force increase and decrease protocol that results in protein unfolding and refolding, respectively. Each domain unfolding can be detected as a step-wise increase in bead height.

Using the experimental protocol detailed in the previous paragraph, the number of protein domains that were refolded (the first cycle at each condition is omitted) can be obtained by counting the number of stepwise bead height increase, which corresponds to the unfolding of one I27 protein domain. The bead height increase *ΔH(f)* depends on force and is equivalent to the extension difference of unfolded I27 and folded I27, Δ*x(f)* = *x*_unfolded_(*f*) – *x*_folded_(*f*). Δ*x(f)* translates into tether contour length increase of Δ*L*_0_ by the Marko-Siggia formula [41]:

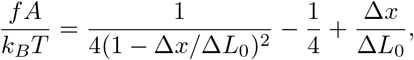

where *A* ∼ 1 nm is the persistence of the unfolded polyprotein of I27 [32]. Multiple cycles of (I27)_8_ folding-unfolding yields Δ*L*_0_ = 27 ± 5 nm (data not shown). A home-written MATLAB code was used for semiautomatic counting of the unfolding steps, with Δ*L*_0_ in the range of 27 ± 2σ nm set as the criteria for assigning the apparent bead height increase as protein unfolding. Based on this method, the fraction of refolded (I27)_8_ at each cycle can be obtained.

## RESULTS

### DnaK in the apo– and ADP-bound state is an effective holdase

We used single protein refolding assay as detailed in the Methods section to probe the action of DnaK in regulating the refolding of (I27)_8_ tether. In this assay, the number of step-wise increase in bead height corresponds to the number of protein domains that are unfolded, which gives us the number of protein domains that are refolded in the preceding cycle (Fig. 1). The folding rate of each I27 domain in the absence of binding partner, *k*_f_(*f*), is ∼ 10 s^-1^ at 1 pN [28]. In the absence of binding partner, we found that the fraction of (I27)_8_ srefolding is 0.88 ± 0.12 (Fig 2), indicating that the majority of the I27 domains were refolded in the time window of the force cycle protocol that we use in our experiments (see Methods for details). The fractions of protein refolding were obtained from multiple folding-unfolding cycles of (I27)_8_ obtained from multiple independent tethers, and the error bars represent the standard deviation from multiple cycles.

**FIG. 2.**
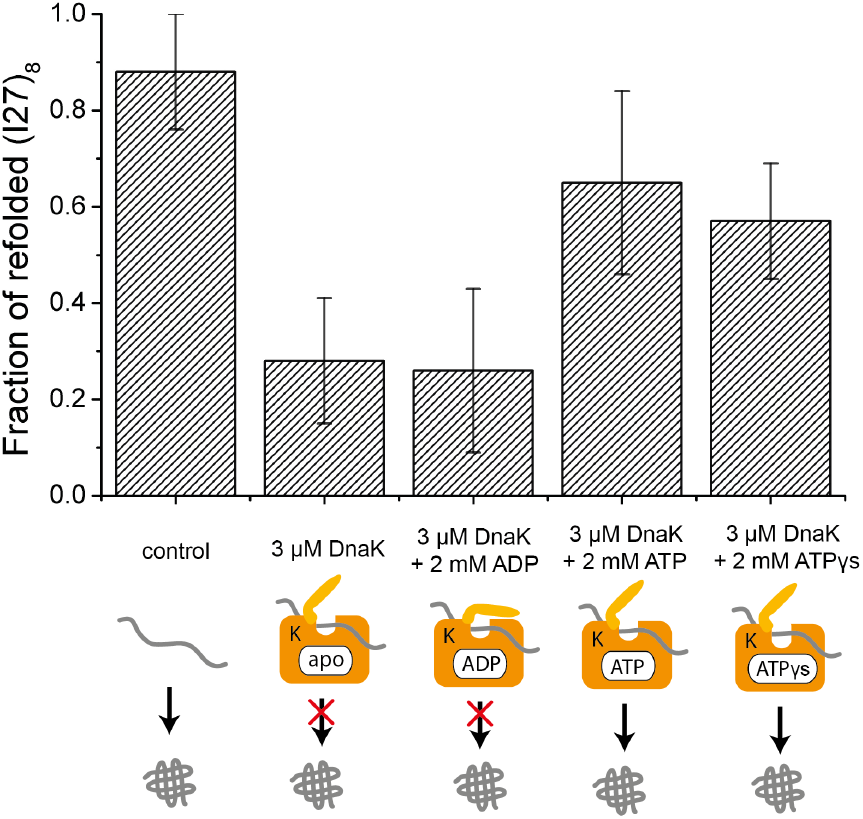
DnaK blocks refolding in the apo– and ADP-state, and allows refolding upon ATP binding. Addition of DnaK in the apo– and ADP-state results in a significantly lower refolding fraction (0.28 ± 0.13 and 0.26 ± 0.17, respectively) compared to the control experiment, indicating the protein being held in the unfolded state. This holding can be relieved by ATP or ATP-*γ*-S binding to DnaK, which results in higher fraction of refolded protein domains.

It is well known that DnaK undergoes conformational changes induced by nucleotide binding [8, 42], which in turn dictates the substrate binding and release activity. In the ATP bound state, DnaK predominantly adopts a more “open” conformation, while in the ADP-state DnaK predominantly adopts a more “closed” conformation. While the conformational state of DnaK cannot be inferred in our single-protein refolding assay, the substrate binding activity can be directly measured. Following force-increase scan and subsequent unfolding of all protein domains in the tether, chaperone holding on to the unfolded (I27)_8_ polyprotein is expected to reduce the fraction of domain refolding at 1 ± 0.1 pN within the 10 s holding time. We first tested DnaK in the absence of nucleotide in solution. We quantified the combined amount of ATP and ADP in our protein stock to be <1% (see Methods) so the majority of DnaK is in apo– (nucleotide-free) state. We found that the fraction of refolding in the presence of 3 *μ*M DnaK is significantly decreased to 0.28 ± 0.13. We did not find conditions in which DnaK completely inhibits protein refolding, possibly due to the dynamic nature of the interaction. Whether DnaK in the apo-state plays a major role *in vivo* is not well understood.

We also measured the fraction of refolded proteins obtained in the presence of 3 *μ*M DnaK and 2 mM ADP, which has been purified from ATP contamination (with ATP contamination <0.01%, see Methods). The fraction of refolded protein in the presence of ADP is similar to the fraction found for DnaK in the apo-state (0.26 ± 0.17). The low fraction is expected as DnaK in the ADP-bound state has a high affinity to peptide substrate, with the SBD-*α* lid closing against the substrate binding groove in SBD-*β* [5]. Therefore, DnaK in the ADP-bound state also functions as a holdase.

### ATP binding relieves DnaK’s holdase activity

Next, we investigate DnaK’s activity in the presence of 3 *μ*M DnaK and 2 mM ATP (Fig. 3). We found that the fraction of refolded protein domains (0.65 ± 0.19) is much higher as compared to those in apo-state. This is consistent with the low substrate binding affinity and “open” conformation of DnaK in the ATP-bound state, which can relieve the holdase activity of DnaK. This fraction, however, is less as compared to those obtained in the absence of chaperone.

**FIG. 3.**
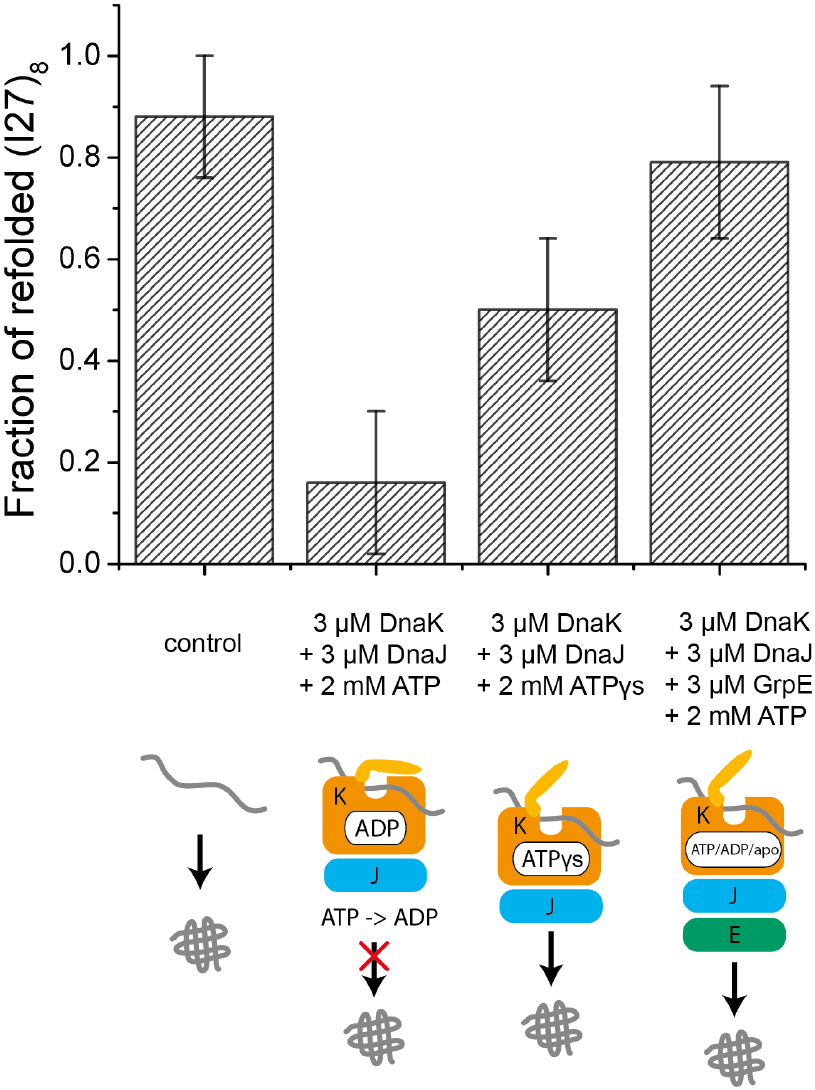
DnaJ and GrpE modulates the DnaK holdase behavior. DnaK/DnaJ blocks protein refolding with addition of 2 mM ATP, but not with ATP-*γ*-S. In the presence of 2mM ATP, the complete DnaK/DnaJ/GrpE system relieves DnaK’s holdase activity, allowing the protein domains to refold.

DnaK in the presence of ATP can assume either an ATP-bound state or ATP can be hydrolyzed into ADP. While ATP hydrolysis rate is very slow in the absence of co-chaperone DnaJ, it has been suggested that substrate binding can induce ATP hydrolysis by 25-fold to allow for spontaneous substrate locking [11–13]. So far, it is not clear whether the protein refolding that we observed in the presence of 2 mM ATP is due to ATP binding or ATP hydrolysis. To distinguish between the two possible scenarios, we performed experiments in the presence of non-hydrolyzable nucleotide analogs ATP*γ*s (2 mM). If ATP hydrolysis is needed for substrate release, we would expect a lower fraction of refolded proteins domains. Our result shows that the refolded domain fraction is 0.57 ± 0.12, which suggests that ATP binding alone is enough to induce DnaK substrate release. In addition, our results also suggest that substrate binding alone is not sufficient to induce ATP hydrolysis, at least in our experimental condition and timescale.

### Co-chaperones DnaJ and GrpE control DnaK’s holdase activity

We next probed protein refolding in the presence of 3 *μ*M DnaK, 3 *μ*M DnaJ, and 2 mM ATP (Fig 3). As DnaJ is expected to significantly accelerate ATP hydrolysis upon DnaK-ATP binding to substrate, we would expect that DnaK assumes a “closed” conformation and decreases the fraction of protein refolding. Indeed, we found that addition of DnaJ significantly decreased the fraction of refolded protein domains to 0.16 ± 0.14, as compared to 0.65 ± 0.19 obtained in the absence of DnaJ. This decrease cannot be explained by DnaJ holding to unfolded substrate, as the fraction of protein refolding obtained in the presence of DnaJ with or without ATP is > 0.5 (Fig 4). Therefore, the significant decrease in fraction of protein refolding is likely caused by DnaJ catalyzing ATP hydrolysis in DnaK. In contrast, we did not observe significant reduction in the fraction of refolded protein domains when we performed similar experiments in the presence of 3 *μ*M DnaK, 3 *μ*M DnaJ, and 2 mM ATP*γ*s. Therefore, substrate (i.e., unfolded protein) release from DnaK is induced by ATP binding, while ATP hydrolysis is responsible for substrate locking to DnaK presumably bound with ADP.

**FIG. 4.**
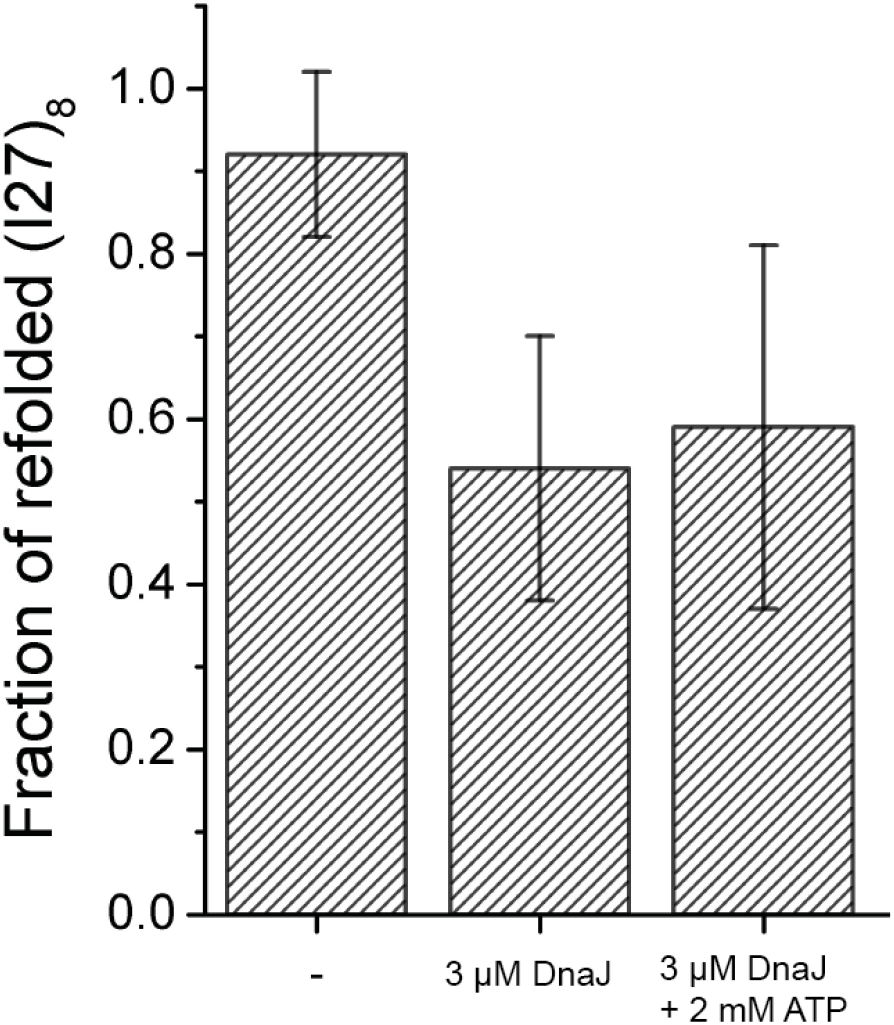
DnaJ functions as a moderate ATP-independent holdase. Addition of DnaJ alone or with 2 mM ATP results in a moderate reduction in the fraction of domains refolded.

Next we investigated protein refolding in the presence of 3 *μ*M DnaK, 3 *μ*M DnaJ, 3 *μ*M GrpE, and 2 mM ATP, which forms a complete chaperone machinery in *E. coli*. GrpE is a nucleotide exchange factor (NEF) that facilitates ADP release from DnaK. Together with DnaJ and GrpE, the nucleotide binding state of DnaK can cycle between multiple states, with different substrate binding activities as characterized previously. The fraction of refolding is relatively high (0.79 ± 0.15), indicating that holdase activity of DnaK is minimal in this condition. This result is consistent with the nucleotide exchange role of GrpE which allows DnaK to switch to the ATP-bound form.

## DISCUSSION

In this article, we investigated the molecular mechanism of *E. Coli* Hsp70 (DnaK) on a single-molecule level using magnetic tweezers. Previously, this method has been used to study trigger factor chaperone, which was shown to work as a foldase [43]. We discovered that DnaK can function as a holdase, depending on the nucleotide binding state. It acts as a strong holdase in the apo– and ADP bound state, which is released in the ATP-bound state. The balance between these states is regulated by its co-chaperones DnaJ and GrpE through catalyzing hydrolysis of ATP and release of ADP.

Our results suggest that DnaK can kinetically stabilize the unfolded state or polypeptide chain in the apo-state, ADP-bound state, or in the presence of DnaJ and ATP. This is shown by the significant decrease in the fraction of protein refolding under these conditions (Fig 2). It is interesting that DnaK in the apo-state can function as a very effective holdase. Hsp70 family of proteins is often thought to be in mainly nucleotide-bound form (either ATP– or ADP-bound). However, recent spFRET experiments on Ssc1, a yeast mitochondrial Hsp70 that bears >50 % sequence identity to DnaK, revealed a frequent transition to a nucleotide-free state [44], which highlight the possible importance of this nucleotide-free state in other Hsp70 family members.

We also found that the holdase activity of DnaK is largely released when it is bound with ATP, indicating a weaker binding of DnaK to the unfolded protein. In the presence of cochaperones DnaJ and GrpE, the nucleotide-bound state of DnaK can cycle between ATP-bound, ADP-bound and possibly apo-states, which in turn enables DnaK binding to the unfolded protein in cycles of tighter and weaker binding modes. Such a mechanism on one hand slows down the overall folding and on the other hand prevents complete inhibition of refolding. This combined cycle of DnaK’s holding and release activity may result in slower overall folding process. Although slowing down protein folding appears to be counter-intuitive as holdase implies anti-folding, this may actually facilitate the transition from unfolded state into its native folded state by altering the folding pathway on a free energy landscape that potentially reduces the possibility of being trapped in metastable misfolded intermediates. We surmise that the role of chaperone to guide the client protein folding into its native/functional state is more crucial than accelerating protein folding.

Previously, the chaperone activity of DnaK/DnaJ/GrpE system has been investigated with atomic force microscopy (AFM)-based pulling on (I27)_8_ [45]. The authors found that mechanically unfolded I27 domains tend to misfold, as characterized by significantly shorter or longer contour lengths released during the unfolding segment by moving the cantilever away from the surface. They found that the DnaK/DnaJ/GrpE system in the presence of ATP resulted in synergy that suppressed the misfolded states. In our assay, such misfolded states were not observed, which can be caused by several differences between the two experiments: 1) the states of the unfolded polyprotein might be different, as the forces applied to the unfolded polyprotein were below 100 pN in our experiments, while it could reach 300 pN in the AFM experiments. 2) The folding conditions might be different. In our assay, the unfolded I27 polyprotein was refolded at 1 ± 0.1 pN, while in the AFM assay the folding took place after the cantilever reapproached the surface, with some degree of uncertainty in the level of force applied to the tether. 3) The protein constructs might be different between ours and Nunes et al. In wild type I27 domain, there are two cysteines at residues 47 and 63 [34]. To avoid potential formation of disulfide bond between the two cysteines, we substituted the two cysteines with alanines. Although our experimental conditions and model substrate do not allow study of DnaK’s action on misfolded conformation, we provide a comprehensive picture of DnaK’s holdase behavior that complements the previous study.

Another publication has shown that DnaK in the ADP-bound state can directly bind and stabilize protein in both the unfolded and folded state of the substrate, obtained with optical tweezers on four maltose binding proteins (4MBP) substrate [46]. The stabilizing effect on the unfolded state is consistent with the holdase function in this article, but we cannot observe the stabilizing effect on the folded state. This can be explained by the substrate selection, as (I27)_8_ already exhibits a high mechanical stability even in the absence of chaperone. In addition, the authors also found that the complete DnaK/DnaJ/GrpE system in the presence of 1 mM ATP can suppress misfolding and/or aggregation, and promotes refolding. Again, as the model substrate that we chose does not have any apparent misfolding behavior, we cannot confirm whether DnaK can suppress mis-folding and/or aggregation using our model substrate. At the time of writing, another group published a study on DnaK-mediated mechanical folding using AFM-based pulling operating in force-clamp mode on ubiquitin and wild type (I27)_8_ as substrate [47]. Our results are largely similar with regard to DnaK’s holdase activity at different experimental conditions.

Our findings are schematized in Fig. 5 with DnaK-client binding being cycled between holding and release mode depending on the nucleotide binding state, cochaperones DnaJ/GrpE, and potentially other factors not accounted for in this study. We also recognize some limitations in our study. First, the findings derived from the model substrate that we choose, (I27)_8_, may not be generalized to other protein substrate. Also, we do not find any misfolding in our model substrate. On one hand, this allows us to provide deeper insights into DnaK’s holdase activity, but on the other hand we cannot get any information on the interplay between DnaK and protein misfolding. As such, it may be worthwhile to explore DnaK’s binding to different model substrate with different sequence and structure to provide a more complete picture of DnaK’s chaperone activity. In summary, we have provided a comprehensive picture of DnaK’s holdase activity and how multiple factors influence the holding and release of protein substrate. Although we largely focus on one aspect of DnaK, i.e. holdase, the role of DnaK is likely to be multifaceted, including but not limited to holdase, foldase, or unfoldase. Further investigation using a plethora of model substrate with diverse properties may be useful to consolidate our understanding of DnaK’s chaperone activity.

**FIG. 5.**
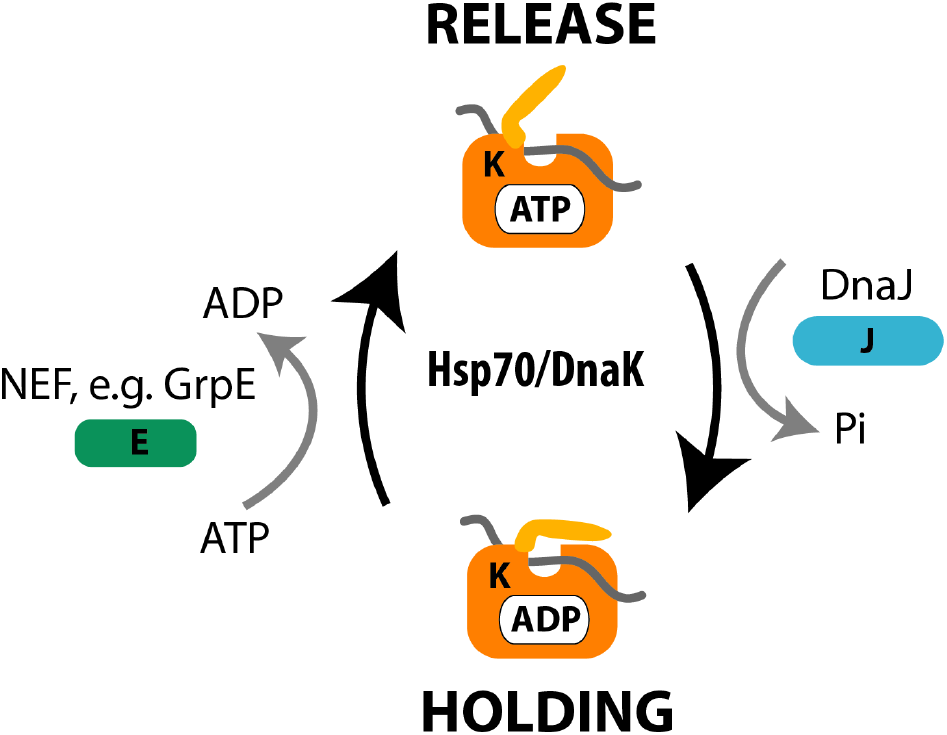
Schematic diagram of DnaK/DnaJ/GrpE chaperone system. DnaK cycles the client protein between holding and release step, which is controlled by its cochaperones DnaJ and GrpE.

## SUPPORTING INFORMATION

*S1 Table*. **Quantification of ATP contamination (in percentage) of the reagents and proteins used in our experiments**. The presence of ATP contamination in the reagents and protein samples used are quantified using a very sensitive luciferase assay (see Materials and methods), to ensure that the observations are not due to contamination of ATP.

## ACKNOWLEDGMENTS

We would like to thank the protein production platform (PPP) in the School of Biological Sciences at Nanyang Technological University for the expression and purification of DnaK, DnaJ, and GrpE.

## References

[1] DA Parsell and Susan Lindquist. The function of heat-shock proteins in stress tolerance: degradation and reactivation of damaged proteins. Annual review of genetics, 27(1):437–496, 1993.

[2] F Ulrich Hartl, Andreas Bracher, and Manajit Hayer-Hartl. Molecular chaperones in protein folding and proteostasis. Nature, 475(7356):324, 2011.

[3] Giuseppina Turturici, Gabriella Sconzo, and Fabiana Geraci. Hsp70 and its molecular role in nervous system diseases. Biochemistry research international, 2011, 2011.

[4] Jason Chung, Anh-Khoi Nguyen, Darren C Henstridge, Anna G Holmes, MH Stanley Chan, Jose L Mesa, Graeme I Lancaster, Robert J Southgate, Clinton R Bruce, Stephen J Duffy, et al. Hsp72 protects against obesity-induced insulin resistance. Proceedings of the National Academy of Sciences, 105(5):1739–1744, 2008.

[5] Eric B Bertelsen, Lyra Chang, Jason E Gestwicki, and Erik RP Zuiderweg. Solution conformation of wild-type e. coli hsp70 (dnak) chaperone complexed with adp and substrate. Proceedings of the National Academy of Sciences, 106(21):8471–8476, 2009.

[6] Ruifeng Qi, Evans Boateng Sarbeng, Qun Liu, Katherine Quynh Le, Xinping Xu, Hongya Xu, Jiao Yang, Jennifer Li Wong, Christina Vorvis, Wayne A Hendrickson, et al. Allosteric opening of the polypeptide-binding site when an hsp70 binds atp. Nature Structural and Molecular Biology, 20(7):900, 2013.

[7] Roman Kityk, Jurgen Kopp, Irmgard Sinning, and Matthias P Mayer. Structure and dynamics of the atp-bound open conformation of hsp70 chaperones. Molecular cell, 48(6):863–874, 2012.

[8] Anastasia Zhuravleva and Lila M Gierasch. Substrate-binding domain conformational dynamics mediate hsp70 allostery. Proceedings of the National Academy of Sciences, 112(22):E2865–E2873, 2015.

[9] Matthias P Mayer. Hsp70 chaperone dynamics and molecular mechanism. Trends in biochemical sciences, 38(10):507–514, 2013.

[10] Wolfgang Rist, Christian Graf, Bernd Bukau, and Matthias P Mayer. Amide hydrogen exchange reveals conformational changes in hsp70 chaperones important for allosteric regulation. Journal of Biological Chemistry, 281(24):16493–16501, 2006.

[11] Rick Russell, A Wali Karzai, Andrew F Mehl, and Roger McMacken. Dnaj dramatically stimulates atp hydrolysis by dnak: insight into targeting of hsp70 proteins to polypeptide substrates. Biochemistry, 38(13):4165–4176, 1999.

[12] Holger Theyssen, Hans-Peter Schuster, Lars Packschies, Bernd Bukau, and Jochen Reinstein. The second step of atp binding to dnak induces peptide release. Journal of molecular biology, 263(5):657–670, 1996.

[13] Robert Jordan and Roger McMacken. Modulation of the atpase activity of the molecular chaperone dnak by peptides and the dnaj and grpe heat shock proteins. Journal of Biological Chemistry, 270(9):4563–4569, 1995.

[14] Andrea D Thompson, Steffen M Bernard, Georgios Skiniotis, and Jason E Gestwicki. Visualization and functional analysis of the oligomeric states of escherichia coli heat shock protein 70 (hsp70/dnak). Cell Stress and Chaperones, 17(3):313–327, 2012.

[15] MP Mayer and B Bukau. Hsp70 chaperones: cellular functions and molecular mechanism. Cellular and molecular life sciences, 62(6):670, 2005.

[16] Rick Russell, Robert Jordan, and Roger McMacken. Kinetic characterization of the atpase cycle of the dnak molecular chaperone. Biochemistry, 37(2):596–607, 1998.

[17] John S McCarty, Alexander Buchberger, Jochen Reinstein, and Bernd Bukau. The role of atp in the functional cycle of the dnak chaperone system. Journal of molecular biology, 249(1):126–137, 1995.

[18] Douglas M Cyr, Thomas Langer, and Michael G Douglas. Dnaj-like proteins: molecular chaperones and specific regulators of hsp70. Trends in biochemical sciences, 19(4):176–181, 1994.

[19] Olivier Genest, Joel R Hoskins, Jodi L Camberg, Shannon M Doyle, and Sue Wickner. Heat shock protein 90 from escherichia coli collaborates with the dnak chaperone system in client protein remodeling. Proceedings of the National Academy of Sciences, 108(20):8206–8211, 2011.

[20] Shannon M Doyle, Shankar Shastry, Andrea N Kravats, Yu-Hsuan Shih, Marika Miot, Joel R Hoskins, George Stan, and Sue Wickner. Interplay between e. coli dnak, clpb and grpe during protein disaggregation. Journal of molecular biology, 427(2):312–327, 2015.

[21] Pierre Goloubinoff, Axel Mogk, Anat Peres Ben Zvi, Toshifumi Tomoyasu, and Bernd Bukau. Sequential mechanism of solubilization and refolding of stable protein aggregates by a bichaperone network. Proceedings of the National Academy of Sciences, 96(24):13732–13737, 1999.

[22] Michal Zolkiewski. Clpb cooperates with dnak, dnaj, and grpe in suppressing protein aggregation a novel multichaperone system from escherichia coli. Journal of Biological Chemistry, 274(40):28083–28086, 1999.

[23] Giulia Calloni, Taotao Chen, Sonya M Schermann, Hung-chun Chang, Pierre Genevaux, Federico Agostini, Gian Gaetano Tartaglia, Manajit Hayer-Hartl, and F Ulrich Hartl. Dnak functions as a central hub in the e. coli chaperone network. Cell reports, 1(3):251–264, 2012.

[24] Sergey V Slepenkov and Stephan N Witt. The unfolding story of the escherichia coli hsp70 dnak: is dnak a holdase or an unfoldase? Molecular microbiology, 45(5):1197–1206, 2002.

[25] Sandeep K Sharma, Philipp Christen, and Pierre Goloubinoff. Disaggregating chaperones: an unfolding story. Current Protein and Peptide Science, 10(5):432–446, 2009.

[26] Hu Chen, Hongxia Fu, Xiaoying Zhu, Peiwen Cong, Fumihiko Nakamura, and Jie Yan. Improved high-force magnetic tweezers for stretching and refolding of proteins and short dna. Biophysical journal, 100(2):517–523, 2011.

[27] Hu Chen, Xiaoying Zhu, Peiwen Cong, Michael P Sheetz, Fumihiko Nakamura, and Jie Yan. Differential mechanical stability of filamin a rod segments. Biophysical journal, 101(5):1231–1237, 2011.

[28] Hu Chen, Guohua Yuan, Ricksen S Winardhi, Mingxi Yao, Ionel Popa, Julio M Fernandez, and Jie Yan. Dynamics of equilibrium folding and unfolding transitions of titin immunoglobulin domain under constant forces. Journal of the American Chemical Society, 137(10):3540–3546, 2015.

[29] Mingxi Yao, Wu Qiu, Ruchuan Liu, Artem K Efremov, Peiwen Cong, Rima Seddiki, Manon Payre, Chwee Teck Lim, Benoit Ladoux, Rene-Marc Mege, et al. Force-dependent conformational switch of *α*-catenin controls vinculin binding. Nature communications, 5:4525, 2014.

[30] Mingxi Yao, Benjamin T Goult, Benjamin Klapholz, Xian Hu, Christopher P Toseland, Yingjian Guo, Peiwen Cong, Michael P Sheetz, and Jie Yan. The mechanical response of talin. Nature communications, 7:11966, 2016.

[31] Shimin Le, Xian Hu, Mingxi Yao, Hu Chen, Miao Yu, Xiaochun Xu, Naotaka Nakazawa, Felix M Margadant, Michael P Sheetz, and Jie Yan. Mechanotransmission and mechanosensing of human alpha-actinin 1. Cell reports, 21(10):2714–2723, 2017.

[32] Ricksen S Winardhi, Qingnan Tang, Jin Chen, Mingxi Yao, and Jie Yan. Probing small molecule binding to unfolded polyprotein based on its elasticity and refolding. Biophysical Journal, 111(11):2349–2357, 2016.

[33] Pavel Savitsky, James Bray, Christopher DO Cooper, Brian D Marsden, Pravin Mahajan, Nicola A Burgess-Brown, and Opher Gileadi. High-throughput production of human proteins for crystallization: the sgc experience. Journal of structural biology, 172(1):3–13, 2010.

[34] Sabina Improta, Anastasia S Politou, and Annalisa Pastore. Immunoglobulin-like modules from titin i-band: extensible components of muscle elasticity. Structure, 4(3) :323–337, 1996.

[35] Sung-Jong Jeon and Kazuhiko Ishikawa. A novel adp-dependent dna ligase from aeropyrum pernix k1. FEBS letters, 550(1-3):69–73, 2003.

[36] Masaru Nakatani, Satoshi Ezaki, Haruyuki Atomi, and Tadayuki Imanaka. A dna ligase from a hyperthermophilic archaeon with unique cofactor specificity. Journal of bacteriology, 182(22):6424–6433, 2000.

[37] Huijuan You, Simon Lattmann, Daniela Rhodes, and Jie Yan. Rhau helicase stabilizes g4 in its nucleotide-free state and destabilizes g4 upon atp hydrolysis. Nucleic acids research, 45(1):206–214, 2016.

[38] Xiaodan Zhao, Xiangjun Zeng, Chen Lu, and Jie Yan. Studying the mechanical responses of proteins using magnetic tweezers. Nanotechnology, 28(41):414002, 2017.

[39] Georgyi V Los, Lance P Encell, Mark G McDougall, Danette D Hartzell, Natasha Karassina, Chad Zimprich, Monika G Wood, Randy Learish, Rachel Friedman Ohana, Marjeta Urh, et al. Halotag: a novel protein labeling technology for cell imaging and protein analysis. ACS chemical biology, 3(6):373–382, 2008.

[40] Bijan Zakeri, Jacob O Fierer, Emrah Celik, Emily C Chittock, Ulrich Schwarz-Linek, Vincent T Moy, and Mark Howarth. Peptide tag forming a rapid covalent bond to a protein, through engineering a bacterial adhesin. Proceedings of the National Academy of Sciences, 109(12):E690–E697, 2012.

[41] John F Marko and Eric D Siggia. Stretching dna. Macromolecules, 28(26):8759–8770, 1995.

[42] Alexander Buchberger, Holger Theyssen, Hartwig Schroder, John S McCarty, Giuseppe Virgallita, Philipp Milkereit, Jochen Reinstein, and Bernd Bukau. Nucleotide-induced conformational changes in the atpase and substrate binding domains of the dnak chaperone provide evidence for interdomain communication. Journal of Biological Chemistry, 270(28):16903–16910, 1995.

[43] Shubhasis Haldar, Rafael Tapia-Rojo, Edward C Eckels, Jessica Valle-Orero, and Julio M Fernandez. Trigger factor chaperone acts as a mechanical foldase. Nature communications, 8(1):668, 2017.

[44] Martin Sikor, Koyeli Mapa, Lena Voith von Voithenberg, Dejana Mokranjac, and Don C Lamb. Real-time observation of the conformational dynamics of mitochondrial hsp70 by spfret. The EMBO journal, 32(11):1639–1649, 2013.

[45] João M Nunes, Manajit Mayer-Hartl, F Ulrich Hartl, and Daniel J Muller. Action of the hsp70 chaperone system observed with single proteins. Nature communications, 6:6307, 2015.

[46] Alireza Mashaghi, Sergey Bezrukavnikov, David P Minde, Anne S Wentink, Roman Kityk, Beate Zachmann-Brand, Matthias P Mayer, Günter Kramer, Bernd Bukau, and Sander J Tans. Alternative modes of client binding enable functional plasticity of hsp70. Nature, 539(7629):448, 2016.

[47] Judit Perales-Calvo, David Giganti, Guillaume Stirnemann, and Sergi Garcia-Manyes. The force-dependent mechanism of dnak-mediated mechanical folding. Science Advances, 4(2):eaaq0243, 2018.

